# Genomic Hotspots: Localized chromosome gene expansions identify lineage-specific innovations as targets for functional biodiversity and predictions of stress resilience

**DOI:** 10.1101/2024.05.23.594666

**Authors:** Eric Edsinger, Leonid L. Moroz

## Abstract

Functional and biodiversity genomics is essential for assessment and monitoring of planetary health and species-specific management in changing ecosystems. However, experimental knowledge of gene functions is limited to a few species, and dependencies on distantly related models. Combined with unrecognized degrees of lineage-specific gene family expansion, this means that traditional comparative methods are insufficient. Here, we clarify definitions of homology and genomic ‘dark matter’ and introduce the concept of a hotspot, defined as innovations underlying the evolution of lineage-specific biology. We illustrate hotspots using molluscs having chromosome-scale genome assemblies and focus on heat-sensing TRPM channels and species living in environments of extreme heat stress (e.g., high intertidal and hydrothermal vent gastropods and bivalves). Integrating gene family, orthogroup, and domain-based methods with genomic hotspots (local paralog expansions on chromosomes), we show that conventional approaches overlook substantial amounts of species-specific gene family diversity due to limitations of distant homology detection. In contrast, local segmental duplications are often recent, lineage-specific genetic innovations reflecting emerging adaptions and can be identified for any genome. Revealed TRPM gene family diversification highlights unique neural and behavioral mechanisms that could be beneficial in predicting species’ resilience to heat stress. In summary, the identification of hotspots and their integration with other types of analyses illuminate evolutionary (neuro)genomic strategies that do not depend on knowledge from model organisms and unbiasedly reveal evolutionarily recent lineage-specific adaptations. This strategy enables discoveries of biological innovations across species as prospective targets for modeling, management, and biodiversity conservation.

## 2 Introduction

Environmental impacts, including record-setting marine heat waves(1,2), are affecting global biodiversity and planetary health(2–6). For marine ecosystems, recovery may be slow due to the massive heat-buffer capacity of oceans(6–10), but understanding how local species respond to accelerating environmental extremes is critical to biodiversity management. For example, Marine Protected Areas are isolated refugia with connectivity for benthic marine invertebrates provided by recruitment of pelagic swimming larvae(11–14). However, a larva’s binary decision to undergo settlement or not can be temperature sensitive (15–17), with implications for species survivorship and distribution in management. Powerful, accessible approaches to predict the adaptive potential of local species are needed for long-term modeling and mitigation of environmental impacts.

Biodiversity Genomics and the umbrella Earth BioGenome Project aim to produce reference genomes with chromosome assemblies for every eukaryotic species(18–25), including diverse spiralians(26,27), with opportunities to address environmental stresses(19,21–25). Yet, genomic data are generally not accompanied by molecular-functional knowledge. Furthermore, there are limited tools to evaluate adaptive potential from diverse lineages(20–22,25,28,29), and integrative approaches across fields, like neuroscience and conservation biology(20,25,28,30–32).

Homology-based annotation of gene function is commonly used in the absence of direct molecular knowledge, wherein sequence and increasingly structural similarities enable mapping of gene function in genetic models, like humans, *Drosophila melanogaster*, and *Caenorhabditis elegans*, to a target species(33–39). Still, gene families present in a target species but absent in a popular reference species can go mis-annotated or unannotated. Similarly, detection of the phylogenetic signal of remote homologs in sequence alignment becomes difficult at 20-35% sequence similarity (twilight zone) and goes beyond the theoretical limits at less than 20% (midnight zone)(40–43), though deep-learning structural approaches are pushing these limits(33–35,39,44). For instance, it is common for 25% or more of genes in a spiralian genome to go unannotated.

Genetic innovations underlying speciation adaptations can be most relevant to functional biodiversity assessments(45–52) but are the most likely to go undetected in current bioinformatic pipelines (detailed below; Figure 1A)(37,53). Even the well-studied *Drosophila* has over 500 unannotated genes that arose during the recent evolution of its genus(53). Overall, annotation methods that reduce reliance on distantly related species and highlight genetic innovations underlying lineage-specific biology are desirable.

**Figure 1.**
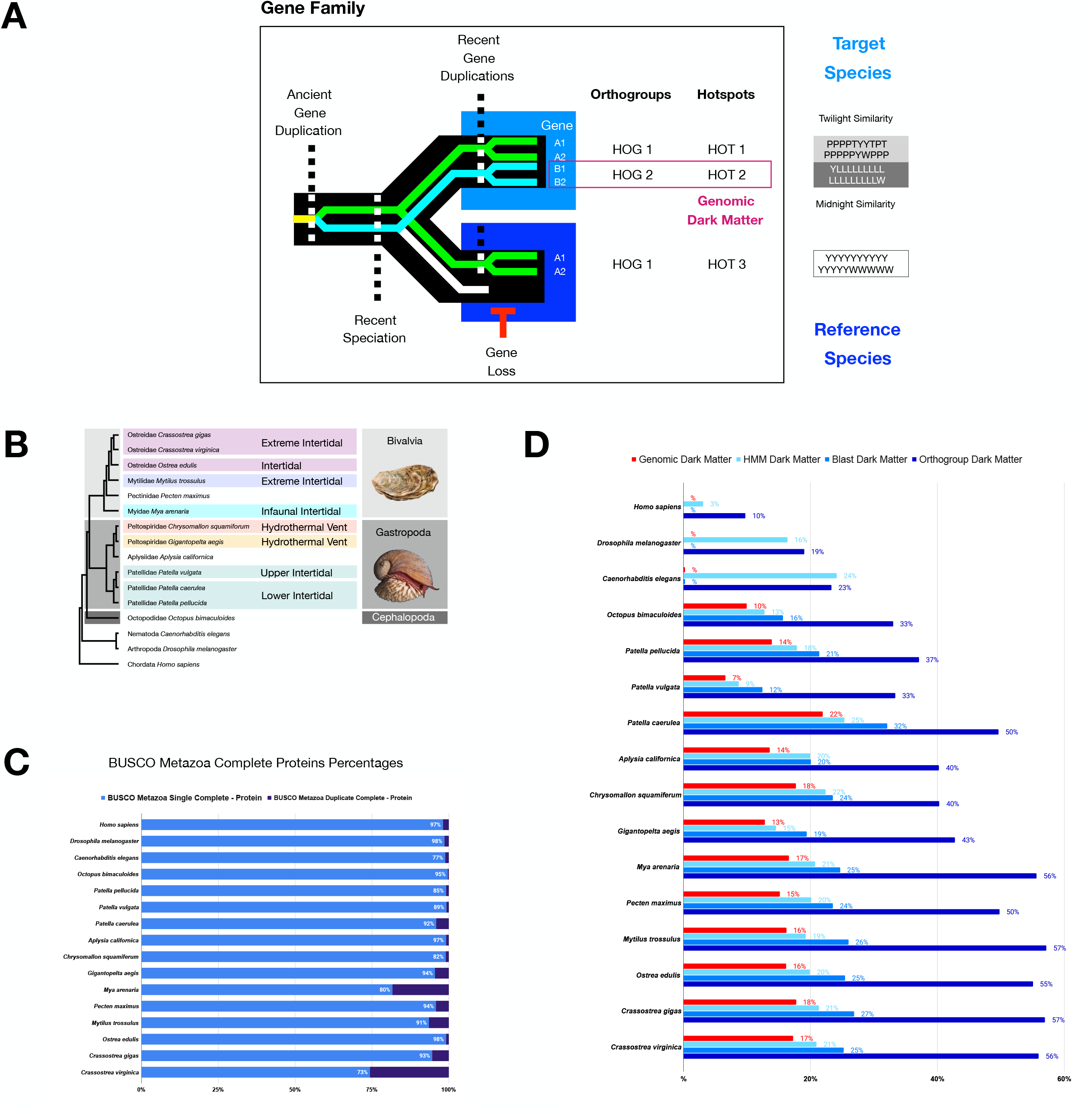
Genomic dark matter, Species16 species, and their proteomes. 1A) A schematic highlighting one possible scenario where genomic dark matter arises due to patterns of gene and species evolution. 1B) The phylogenetic tree for species and their habitats highlighting thermal stress. Molluscan classes are indicated in color blocks with cephalopods dark gray, gastropods medium gray, and bivalves light gray. Independent origins of heat-stress habitats (intertidal or hydrothermal vent) are indicated in color blocks. Four independent origins of intertidal habitats or upper-intertidal regions include 1) oyster - purple, 2) mussel - light purple, 3) *Mya* - light blue, and 4) snail - teal. Two independent origins of hydrothermal vent habitats are 1) *Chrysomallon* light red and 2) *Gigantopelta* light orange. 1C) BUSCO Metazoa genome assessments for completeness and duplication. 1D) Percentage of genomic dark matter in each species, where genomic dark matter is defined as genes that lacked functional annotation based on sequence homology to reference species and their functionally assessed genes. Assessments for functional annotations were based on 1) HMM-based GO-Pfam domain and PANTHER gene family annotations, 2) top hit in reference genomes based on one-direct Blast annotations, and 3) Diamond-based genome clustering of all Species16 species annotations (see Methods for details).

In deciphering those mechanisms, it was established that gene families commonly expand through segmental duplication of chromosome regions during DNA replication, generating new gene copies (paralogs) physically adjacent to existing gene copies on a chromosome(54–57). Evolutionary recent gene copies can diverge in function (division of labor), enabling biological novelties(54,58,59). Parallel processes, including deletions, inversions, and translocations, result in the spatial mixing of genes, with chromosomes forming “bags of genes” over time(57,60–66). Thus, initial clusters of newly formed gene copies will eventually disperse in the evolution of genomes, meaning localized gene copies on a chromosome likely reflect recent evolutionary events and underlie lineage-specific biology. These genomic regions could also act as catalytic sources of molecular innovations, taking advantage of the proximity of genes and their regulatory sites.

We introduce the concept of hotspots and focus on the genome. Genomic hotspots can be identified using simple bioinformatic methods for reference-free identification of gene copies locally clustered in a genome and integrated with genome-scale homology-based methods of molecular function gene annotation, orthogroups, and gene trees. We focus on genome-sequenced molluscs living in typical vs. extreme heat-stress environments. The selected molluscs include bivalves and gastropods species found in subtidal vs. intertidal or hydrothermal vent habitats. We also epmhasize the Transient Receptor Potential (TRP) ion channel superfamily, including the thermo-sensitive TRPM family(67–70), which are diverse and expressed under heat stress in species used here(71–79). Our findings illustrate that approaches leveraging hotspots could enable predictions of adaptation and resilience in response to environmental change.

## 3 Materials and methods

Genome sources and computational methods are provided as supplementary materials. Python-Unix pipelines are provided as GitHub repository Hotspots_Paper_2024 v0.1.0-alpha.1 (GitHub Hotspots_Paper_2024 v0.1.0-alpha.1: https://github.com/000generic/Hotspots_Paper_2024/tree/v0.1.0-alpha.1). The repository is archived with a permanent DOI at Zenodo (Zenodo DOI: 10.5281/zenodo.11069191: https://zenodo.org/records/11069191).

## 4 Results

### 4.1 Hotspots highlight innovations underlying the origins and evolution of lineage-specific biology

Homology and the origins of novelties are at the core of evolutionary paradigms. Like others, we define homology as a state shared between biological features originating from a feature in a common ancestor. We also expand the definition of genomic dark matter to incorporate genomic structures resistant to functional annotation, especially based on sequence homology (Figure 1A). Finally, we introduce the concept of a hotspot and focus on its application in evolutionary genomics (Figure 1A).

We define a hotspot as the set of innovations underlying the evolution of lineage-specific biology (Figure 1A). Hotspots can be composed of structural components within and across hierarchical levels, from base elements to ecosystems. The term is scaleless. It can have diverse complex contexts, from molecular (e.g., genomic hotspots below) to cellular (e.g., neural circuit hotspots) to organismal (population hotspots) and can include their cross-level integration.

Here we focus on genomic hotspots formed as regions of chromosomes delineated by spatial clusters of gene paralogs. This is similar to synteny, in that the identity of genes in genomic proximity on a chromosome is evaluated, but is distinct, as syntenic methods are defined by identifying patterns across species while hotspots are defined internal to the target without outside reference.

Methodologically, genomic hotspots are free of external requirements of high-quality genome assemblies and annotations outside a given target species or lineage, in contrast to syntenic approaches. Thus, although additional reference genomes can be useful in evaluating hotspots, they are not required in the identification and initial use of hotspots to guide deciphering of novelties and adaptions underlying lineage or species-specific biology and evolution.

To illustrate the ‘hotspot’ approach, we selected 16 genomes with chromosome-level assemblies. Initial assessments of assembly completeness were based on BUSCO Metazoa evaluation of T1 proteomes having one representative sequence per gene, with most species found to be 95% BUSCO Complete or better, but with exceptions of highly derived *Caenorhabditis* (75%), *Patella vulgata* (89%), *Patella pellucida* (86%) and *Chrysomallon squamiformis* (83%). These results are indicative of high-quality genome assemblies and structural gene model annotations (Figure 1C). Additional details are provided in Supplementary Materials.

### 4.2 Genomic dark matter is prevalent in functional biodiversity annotations

We found substantial amounts of genomic dark matter in species genomes after running commonly used functional annotation methods, highlighting limitations of these methods. To illustrate this, we performed three independent types of annotation for the T1 proteomes, specifically: 1) by blasting against best-annotated reference genomes of model organisms (human, *Drosophila*, and *Caenorhabditis*) for one-direction top hit annotations, 2) by blast-based genome clustering for orthogroup annotations, and 3) by HMM-based identification of gene features for protein domain and gene family annotations. We then determined what percentage of genes were annotated or not for each method and across all methods, with genes going undetected in all three declared genomic dark matter.

For Blast annotations, unannotated genes ranged from 12-32% of the genome, with a 23% average (SD 5%) (Figure 1D). The method was intermediate in its ability to annotate but common e-value cut-offs of 1e-3 to 1e-10 mean there can be promiscuous domains and low-level false positives complicating the annotations in unknown ways.

For orthogroup annotations, we found unannotated genes ranged from 33-57% of the genome with a 47% average (SD 9) (Figure 1D). This method is the most powerful for inference of gene function, as its scope of comparison is restricted to orthogroup orthologs across species, thereby avoiding most false-positive issues. However, it is also the most conservative approach, producing exceedingly high levels of unannotated genes, more than double the other two methods, and lacking identification of deeper levels of homology commonly of interest.

For HMM-based domain annotation using Pfam and GO and HMM-based gene family assessment using PANTHER, unannotated genes ranged from 9-25% of the genome with a 19% average (SD 4) (Figure 1D). Although some degree of misannotation due to false positives is likely, it is thought that the highly sensitive information-rich aspects of how HMMs are built can reduce this issue in comparison to Blast and other tools (80). Thus, the HMM method is the most effective for functional gene annotation in Species16 biodiversity genomes but still leaves significant numbers of genes unannotated.

Finally, we find that 7-22% of genes in a genome remained unannotated across all three methods, forming conservatively defined genomic dark matter (Figure 1D). These results highlight the degree to which reference-based methods for the functional annotation of genomes can fail in biodiversity assessments and illustrate the extent to which genetic novelty arises in evolution. We also found that for some species within a genus, their genomes exhibited quite different degrees of unannotated genomic dark matter, for example, 7% vs. 14% vs. 22% in the intertidal snails *P*.*vulgata, P*.*pellucida*, and *Patella caerulea*, respectively, suggesting dynamic patterns of gene innovations in recent speciation.

### 4.3 Genomic hotspots are common in genomes

We developed a simple stand-alone / reference-free method to identify genomic hotspots and found they are common in 16 bilaterian genomes, suggesting their identification can enable targeting of genes underlying species or lineage-specific biology. Focusing on molluscs, we blasted each T1 proteome against itself. We opted for an e-value cutoff of 1e-60 and identified all hits of a query gene located within a window of 20 genes centered on the query gene location in a chromosome or scaffold. These initial clusters were then merged based on linkages of overlapping membership to form final genomic hotspot gene sets per genome.

We found that the number of hotspots and their genes ranged from 483 with 1,982 genes (average ??? genes per hotspot) in the shallow-water octopus *Octopus bimaculoides* to almost 8x as many in the intertidal infaunal clam *Mya arenaria*, with 3,747 hotspots and 11,982 genes (average ??? genes per hotspot). For initial test cases using nineteen species and focusing on hotspot identification in the sea hare *Aplysia californica*, small numbers of false positives arose at an e-value of 1e-40, most likely due to promiscuous domains or motifs. However, a substantially less restrictive e-value of 1e-10 was required to recover the Hox gene complex, an ancient chromosomal gene copy cluster and the most deeply studied and widely recognized (81–86). Also, while larger initial windows of 200 genes, rather than 20, sometimes detected additional hotspot members, the hits often appeared to be distantly related or due to a domain shared between unrelated gene families, greatly increasing false positives. At the same time windows smaller than 20 genes often lost likely hotspot members. Thus, we optimized for a window of 20 genes, as it best provided a stable core number of hotspot true positives, no obvious false positives, and reduced over-aggregation of distantly related hotspots.

### 4.4 Genomic hotspots are enriched in the TRP superfamily and TRPM family

To explore genomic hotspots in the context of gene family evolution, we focused on the TRP superfamily of ion channels and TRPM family within it. We identified all TRP superfamily members for 13 target mollusc species and 3 reference species (Species16). We identified in each species by reciprocal blast, using as queries a reference gene set of all human, *Drosophila*, and *Caenorhabditis* TRP proteins and then blasting back all target hit sequences against the reference proteomes. All target genes having a top hit back to a TRP family member in at least one reference proteome were accepted as candidate homologs. While TRP family size is 17, 22, and 32 genes respectively in *Drosophila, Caenorhabditis*, and human, it varied from 31 in *Octopus* to 167 in the upper intertidal mussel *Mytilus trossulus*, with an average of 81 genes per species (SD 35).

Next, we tested if the TRP superfamily is enriched for hotspots relative to the genome in general for each Species16 species. We found that while the average background density of genome hotspots per 100 genes varied from 3 in *Octopus*, the lowest of all, up to 10 in *Mya*, the average TRP hotspot density varied from 4 in the lower intertidal limpet *Patella caerulea* to 19 in *Mya*. Overall, the TRP superfamily was enriched for hotspots relative to the genome in nearly all selected species, suggesting that TRPs play important roles lineage-specific adaptations across molluscs.

### 4.5 Recent diversification of TRPs in molluscs

To further explore the role of TRPs in molluscan evolution, we assessed patterns of TRP superfamily evolution in Species16 species. Specifically, we constructed phylogenetic trees for the entire TRP superfamily (Figure 2A and 2B), and TRPM in particular (Figure 2C and 2D). We found that *Mytilus* exhibited the greatest diversification with 167 TRP genes (Figure 2E and 2F), including TRPM (Figure 2G and 2H). We find that the majority of TRP gene diversity across species lies outside reference species subbranches, indicating more recent lineage-specific expansions. We also find that hotspot sequences commonly form their own orthogroups, lacking functional annotation (genomic dark matter). Only rarely do hotspots belong to orthogroups that can be functionally annotated based on inclusion of reference genes (Figure 2B and 2D-2H). Finally, we find that species that have independently evolved to live in extreme heat stress environments, such as those found in the intertidal or at hydrothermal vents (Figure 1B), have independently expanded the thermosensitive TRPM gene family, often extensively and uniquely so within the TRP superfamily (Figure 2C, D, G, and H). In bivalves, this includes TRPM gene family expansions within each heat-stress tolerant lineage. In the lineage of oysters, *Ostrea* is lower intertidal and exhibits fewer expansions than the *Crassostrea* species *C. gigas* and *C. virginica*, which are found in the more extreme upper intertidal. Similarly, the mussel *Mytilus* and the clam *Mya* live in heat stress environments of the upper intertidal and infaunal intertidal mudflats, respectively, and exhibit a number of extensive lineage-specific TRPM gene family expansions. In contrast, the scallop *Pecten maximus* is closely related to oysters and mussels but is a subtidal species and has few TRPM genes and no substantial expansions in TRPM diversity (Figure 2C and 2D). In gastropods, the three *Patella* species are intertidal vs.

**Figure 2.**
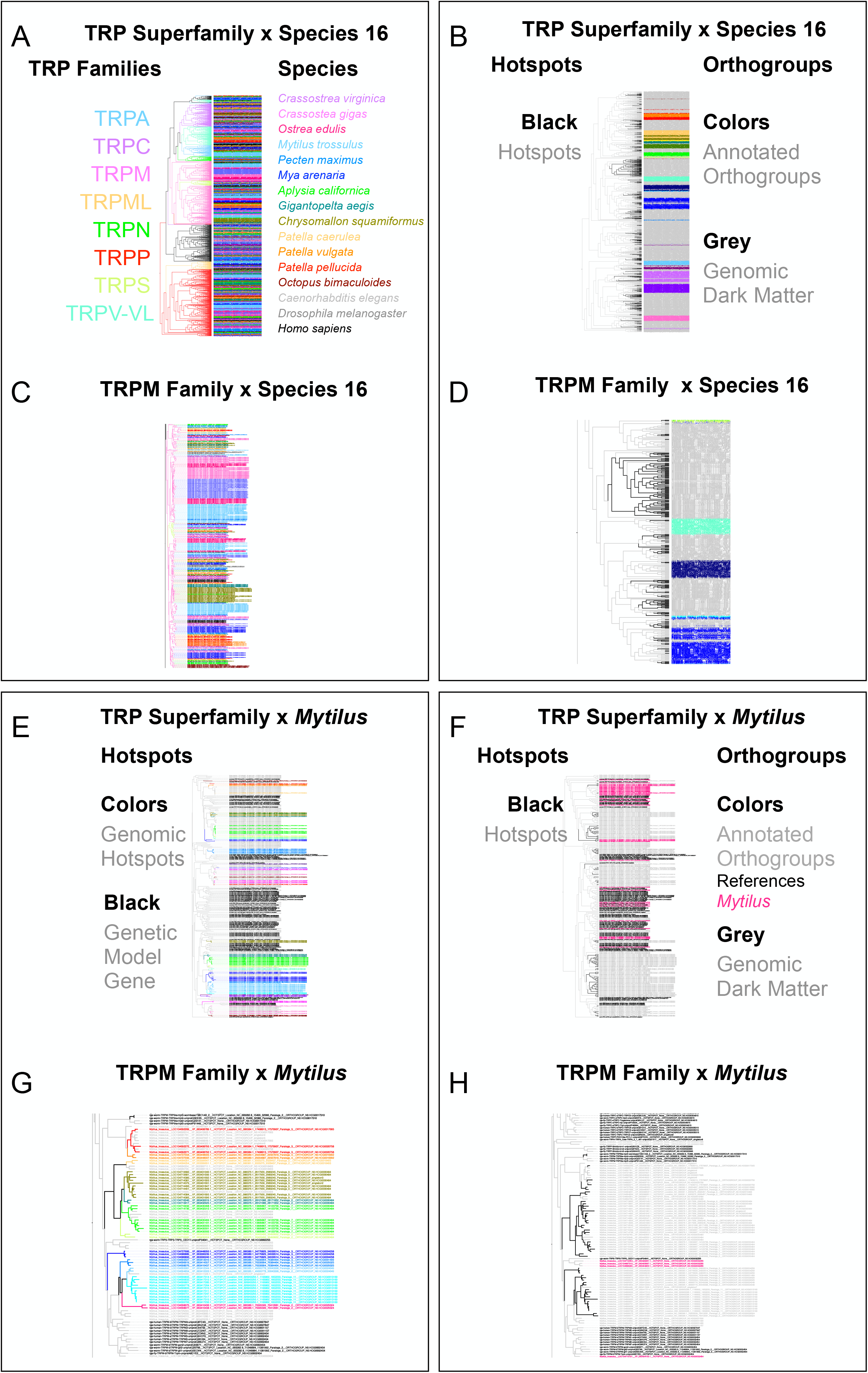
Integration of gene trees, orthogroups, and hotspots for the TRP superfamily in Species 16 species. TRP superfamily and TRPM family (Figure 2A-2D). These evolutionary expansions are visually evident in the gene trees as blocks of species-specific color (Figure 2A and 2B). The color blocks are indicative of multiple paralogous gene copies in a species that arose within its immediate lineage on the Species16 species tree (Figure 1B). For a limited number of subbranches, we observed formation of many fine resolution rainbows of color, indicative of deeply conserved sequences with little lineage-specific evolution since the common ancestor (Figure 2A and 2C). 2A-2H) Integration of gene families, hotspots, and/or orthogroups for TRPs or TRPM and either all Species16 species or Mytilus. 2A) Species16 TRP gene tree with blocks of color on the species side indicative of lineage-specific gene family expansions. 2B) Species16 integration of gene families, hotspots, and orthogroups on the TRP gene tree. Annotated orthogroups are orthogroups that include membership of at least one reference species gene. Genomic dark matter is all sequences not part of an annotated orthogroup. The tree is the same as in panel 2A. 2C) A Species16 TRPM gene tree with blocks of color on the species side indicative of lineage-specific gene family expansions. 2D) Species16 TRPM integration of gene families, hotspots, and orthogroups. The tree is the same as in panel 2C. 2E) Integration of gene families, hotspots, and orthogroups on the TRP gene tree for Mytilus, highlighting individual hotspots in color. 2F) Integration of gene families, hotspots, and orthogroups on the TRP gene tree for Mytilus, highlighting general patterns of hotspots vs orthogroups. The tree is the same as in panel 2E. 2G) Integration of gene families, hotspots, and orthogroups on the TRPM gene tree for Mytilus, highlighting individual hotspots in color. 2H) Integration of gene families, hotspots, and orthogroups on the TRPM gene tree for Mytilus, highlighting general patterns of hotspots vs orthogroups. The tree is the same as in panel 2G.

*Aplysia*, which is a primarily shallow-water subtidal species. Although patterns of gene expansion are less striking, the *Patella* species have more TRPM genes than *Aplysia* and with more small-scale expansions of 1 or 2 genes (Figure 2C and 2D). The two hydrothermal vent gastropods, *Chrysomallon squamiferum* and *Gigantopelta aegis*, belong to the same family but they have adapted to the extreme heat stress independently(77–79). Their genomes show striking patterns of parallel expansion in TRPM genes, less so than expansions found in bivalves but much greater than expansions seen in the other gastropods and octopus (Figure 2C and 2D). Interestingly, the two main expansions in each species occur on the same branch within the greater TRPM gene tree (Figure 2C and 2D).

## 5 Discussion

The presented discussions of hotspots, homology and genomic dark matter agree with previous work (90–98) but can help frame comparative genome–scale studies across biodiversity.

First, identification of genomic hotspots provides a reference-free means to identify candidate genes underlying the origins of lineage-specific biology, as their localization represents more recent evolutionary events due to the eventual dispersal of localized genes in eukaryotic genomes, with some notable exceptions, like Hox genes (61,62,64,66,99). It is striking that Species16 hotspots are predominately genomic dark matter and only more rarely associated with orthogroups and gene tree branches that include reference sequences of three model organisms used here. Future genome-scale statistical analyses and modeling could elucidate potential protection of older gene copies from hotspot formation and/or preferential utilization of new copies in more recent lineage-specific biology, perhaps due to associated gene regulatory elements that might be fully intact in older copies but variable in younger ones.

Second, the observed patterns of TRP family evolution are similar to previous studies in molluscs, including oysters (68,70,71,73,74). TRP gene family members form hotspots at substantially greater levels than observed background levels per genome. The TRP hotspots are predominately composed of unannotated genomic dark matter, which highlight the potential roles of TRP ion channels in lineage-specific biology. The number of TRP ion channels in bivalves and gastropods, with relatively simple nervous systems and behaviors, is much greater than that of humans and *Octopus*, which have independently evolved large brains (Moroz, 2009) and sophisticated behaviors suggesting functional pressures that limit gene diversification in complex nervous systems and/or lead to molecular expansion in simpler nervous systems.

The TRPM gene family is recognized as a primary molecular sensor of temperature, including their elevated expression in response to heat stress in oysters and other molluscs (68,70,71,73,74). Upper intertidal and hydrothermal vents are both environments featuring heat extremes, and bivalve and gastropod lineages have independently entered these environments with greatly expanded TRPM gene family diversity through lineage-specific hotspots.

In summary, Identification of the environmental molecular sensors of direct interest as part of newly emerging mechanistic work in functional biodiversity, enable new tools and resources to predict resilience and adaptability of a species facing rapid environmental change.

## 6 Conclusion

Overall, our findings highlight the idea that genomic hotspots represent relatively recent genetic innovations and that their unbiased reference-free identification can provide a novel and potentially powerful means to elucidate genomic mechanisms of evolution and the origins of genes underlying lineage-specific biology without any greater knowledge beyond the genome itself. TRP ion channels are important targets for understanding lineage-specific adaptations under regimes of environmental change and predicting outcomes for populations in response to such impacts.

## 7 Acknowledgements

We warmly thank J. Hsiao for initial design and coding to characterize and automate homology neighborhoods (now genomic hotspots), and gene family phylogenetic trees, working with EE in the Chalasani Lab at the Salk Institute as a part of the GIGANTIC Project. We also wish to thank S. Chalasani and lab for their encouragement and support.

## 8 Contributions

EE conceived hotspots, formulated definitions, designed, coded, ran, and analyzed bioinformatic experiments, wrote an initial draft of the manuscript. EE and LLM discussed key stages of the work, including neurobiology of stress, and jointly edited the manuscript for submission. LLM provided funding and resources.

